# Fast and adaptive protein structure representations for machine learning

**DOI:** 10.1101/2021.04.07.438777

**Authors:** Janani Durairaj, Mehmet Akdel, Dick de Ridder, Aalt DJ van Dijk

## Abstract

The growing prevalence and popularity of protein structure data, both experimental and computationally modelled, necessitates fast tools and algorithms to enable exploratory and interpretable structure-based machine learning. Alignment-free approaches have been developed for divergent proteins, but proteins sharing functional and structural similarity are often better understood via structural alignment, which has typically been too computationally expensive for larger datasets. Here, we introduce the concept of rotation-invariant shape-mers to multiple structure alignment, creating a structure aligner that scales well with the number of proteins and allows for aligning over a thousand structures in 20 minutes. We demonstrate how alignment-free shape-mer counts and aligned structural features, when used in machine learning tasks, can adapt to different levels of functional hierarchy in protein kinases, pinpointing residues and structural fragments that play a role in catalytic activity.

## 1 Introduction

The dual effect of the ever-growing number of protein structures deposited in the Protein Data Bank (Sussman *et al.*, 1998) and dramatically improved protein structure modelling (Senior *et al.*, 2020) has led to an increasing number of studies incorporating structure information for predicting and understanding protein function. Structures are essential to our understanding of protein biology as their form dictates function and they evolve more slowly than sequences. Research questions for which structural data may provide an answer are many and diverse - ranging from searching for remote protein homologs with similar structural folds across the tree of life to exploring the properties of a single protein family in a single species. These two extremes require different approaches, as both the numbers of protein structures involved and the types of insights that can be obtained differ greatly. In the past years, machine learning is proving itself to be crucial in solving these research questions, evident by its meteoric growth in the bioinformatics field. Machine learning algorithms have been applied across divergent protein structures for tasks such as topology classification (Jain *et al.*, 2009), model quality assessment (Cao *et al.*, 2017), ligand pocket prediction (Krivák and Hoksza, 2018), and mutant stability estimation (Berliner *et al.*, 2014). For specific protein families, structure-based machine learning has helped with predicting SH2 domain specificity (Ferraro *et al.*, 2006), modelling the fitness landscape of cytochrome P450s (Romero *et al.*, 2013), finding similarities in telomerases (Lee *et al.*, 2008), and predicting ligand-binding for G-protein coupled receptors (Vass *et al.*, 2016), among others.

Recently, we published Geometricus (Durairaj *et al.*, 2020), a fast alignment-free protein structure embedding approach for describing and comparing divergent proteins. Geometricus defines discrete so-called shape-mers, analogous to sequence *k*-mers, using a set of rotation-invariant moments. The embedding of a protein is then simply the count vector of these shape-mers. This alignment-free approach accurately represents the topological aspects of proteins in machine learning for predicting protein function. The embedding allows for interpretation by mapping predictive shape-mers back to a set of residues in every protein. Given that applications on divergent proteins often encompass tens of thousands of structures, Geometricus provides a good balance between speed and interpretability.

However, for more similar proteins more correspondence between individual residues is expected, and an alignment better captures information about conservation, outliers, and functionally important residue positions. For each residue, a variety of features can be measured according to their relevance to the problem at hand, ranging from amino acid physicochemical properties to electrostatic energies and, in this research, topological properties via rotation-invariant moments. These features when aligned according to a multiple structure alignment generate a matrix that can directly be used as input to a machine learning algorithm. The algorithm looks across the alignment positions for patterns and correlations relevant for predicting the desired response variable. Predictions can be understood in terms of predictive residue positions, which are now easily compared to known catalytic residues or form hypotheses for mutagenesis studies. Gaps in an alignment are considered as missing data, and alignment positions with too many gaps are often discarded, potentially losing out on the predictive power of catalytically important residues split across multiple gap-filled positions. Thus, to generate an alignment-based feature matrix from a set of similar proteins we start with our recently released Caretta multiple structure alignment algorithm, built with the aim of generating high-coverage alignments for use in machine learning (Akdel *et al.*, 2020).

Computational structure modelling, both *de novo* and homology-based, has started to play more of a role in structure-based machine learning research (Berliner *et al.*, 2014; Cavasotto and Palomba, 2015), leading to datasets with up to thousands of protein structures sharing similar functionality and structural folds. These numbers are difficult to handle with current multiple structure alignment approaches which generally scale poorly with the number of proteins (Ma and Wang, 2014). In many cases, this can be attributed to the initial all vs. all pairwise alignment step used to generate a guide tree for subsequent progressive alignment steps. Multiple sequence alignment algorithms such as ClustalW (Thompson *et al.*, 2003), Kalign (Lassmann and Sonnhammer, 2005) and MUSCLE (Edgar, 2004) circumvent this by using alignment-free *k*-tuple similarity, calculated by collecting matching subsequences of length *k* (*k*-mers) from both input sequences, instead of pairwise alignment. This greatly reduces time complexity and allows for near-linear scaling with the number of proteins. With Geometricus this now becomes possible for structure alignment as well, by defining shape-tuple similarity as the collection of matching shape-mers from each protein’s structure, thus completely avoiding the need for all vs. all pairwise structure alignment. The progressive alignment stage still aligns pairs of proteins at a time, and unlike for sequences, the three-dimensional nature of structures necessitates a superposition step in pairwise alignment. This step orients the input protein pair such that distance measures between aligning residues are meaningful. We use rotation-invariant moments to define this initial superposition step. Both of these improvements are incorporated into the Caretta algorithm to give **Caretta-shape**. We demonstrate that Caretta-shape is comparable to other popular structure alignment algorithms in terms of alignment quality and accuracy, while scaling easily to thousands of proteins.

We use the well-studied protein kinase superfamily to demonstrate how Geometricus and Caretta-shape can be used in unison to explore and understand structural similarities and differences between large datasets of protein structures. We showcase both unsupervised and supervised machine learning analyses at the superfamily, family, and subfamily levels, with emphasis on extracting structural insights useful for downstream research.

## 2 Methods

### 2.1 Caretta-shape

We recently released Caretta (Akdel *et al.*, 2020), a software for multiple structure alignment aimed at generating aligned features for use in machine learning algorithms. The advantage of Caretta in the context of machine learning applications lies in its focus on high coverage alignments using a novel consensus weight mechanism, which improves the information content of the aligned features. Here we detail the modifications made to the Caretta algorithm.

#### 2.1.1 Shape-tuple similarity for fast guide tree construction

An all vs. all similarity matrix is constructed for input proteins by calculating the Bray-Curtis similarity between each protein pairs’ Geometricus count vectors (with *k*-mer size *k* = 20 and resolution *m* = 2). The guide tree for determining the order of progressive alignments is constructed using maximum linkage neighbor joining (Saitou and Nei, 1987) on this similarity matrix.

#### 2.1.2 Rotation-invariant moment-based superposition

Caretta-shape replaces the signal- and secondary structure-based superposition scheme of Caretta by moment-based superposition. For each of the two structures to be aligned, four moment invariants are calculated for each residue with a fixed *k*-mer size (set to *k* = 20), 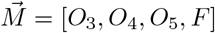 (named as in (Durairaj *et al.*, 2020)). To ensure that the four moments contribute equally to the distance measure, each moment is normalized across both structures by subtracting the mean and dividing by the standard deviation to form 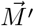. The two series of normalized moment invariants are then aligned by dynamic programming using the Gaussian Caretta score:

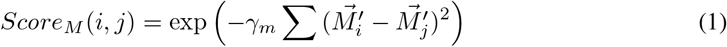

with *γ*_*m*_ = 0.6. The aligning residues are used to calculate the optimal superposition using the Kabsch algorithm (Kabsch, 1976), after which coordinate-based superposition is performed as in the Caretta algorithm with default parameters (*γ* = 0.03, gap open penalty = 1, gap extend penalty = 0.01, consensus weight = 1). Parameter optimization for specific tasks could improve the results presented here, but we leave this open for future exploration.

#### 2.1.3 Benchmarking

Caretta-shape was tested on two benchmark datasets, Homstrad (Mizuguchi *et al.*, 1998) and SABmark-superfamily (Van Walle *et al.*, 2004). The PDB files for these two datasets (390 sets with 3–27 proteins each from Homstrad and 425 sets with 3-42 proteins each from SABmark-superfamily) were obtained from mTM-align’s website (Dong *et al.*, 2017b) and Matt benchmark results (Menke *et al.*, 2008) respectively, in order to directly compare results to the output of these two tools. To this end, the Matt (Menke *et al.*, 2008) and mTM-align (Dong *et al.*, 2017b) alignments for the Homstrad (Mizuguchi *et al.*, 1998) and SABmark-Sup (Van Walle *et al.*, 2004) datasets were obtained from their respective websites. For 17 cases in the Homstrad dataset, mTM-align returned alignments where at least one sequence did not match the corresponding PDB sequence. These cases were not considered.

We report various quality metrics of multiple structure alignments obtained from the different tools benchmarked. For both benchmark datasets we report the average (median) TM-score of the alignment, a measure that takes into account both the structural equivalence of corresponding residues and the overall coverage of the alignment (Zhang and Skolnick, 2004). We also report the median percentage of positions in the alignment without gaps (gapless positions), an aspect important to consider when using aligned features as input to machine learning algorithms, as gaps are seen as missing data and may cause loss of information about the residue positions in which they occur. In addition, the Homstrad dataset provides a set of manually curated reference alignments, for which we define an accuracy score (Acc.) that measures the number of correct gapless positions found, *i.e* gapless positions which are equivalent to positions in the corresponding reference alignment, divided by the total number of gapless positions in the reference alignment.

To estimate Caretta-shape running times, we chose four proteins from the SABMark dataset as “seeds”, with lengths 100, 300, 504, and 714 respectively. Each seed was used to form multiple groups of proteins by introducing noise of up to 5 Å to each of the seed coordinates, to create a given number of members, from 10 to 1010 in increments of 200. Each noisy structure was further rotated by a random angle (between 0 ° and 360 °) along a randomly selected axis. Caretta-shape was then used to align these groups on a Linux workstation using a single thread.

### 2.2 Protein kinases

#### 2.2.1 Data

Protein kinase PDB files with group and family annotations were collected from the kinase–ligand interaction fingerprints and structure database (KLIFS) (Kooistra *et al.*, 2016), resulting in 7,746 monomeric structures collectively named the KinaseAll dataset. Table 1 lists the number of proteins in this dataset in each kinase group.

**Table 1:**
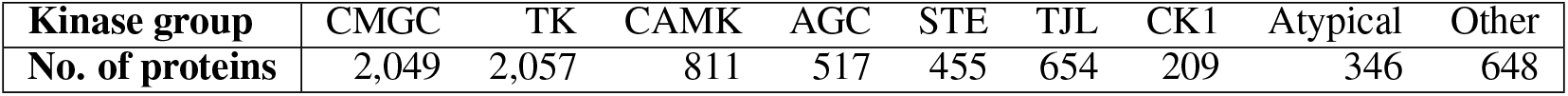
Number of proteins in the KinaseAll dataset across the eight major kinase groups (the rest are labelled as “Other”).

Data about active and inactive states of kinase structures was taken from work by McSkimming *et al.* (2017) yielding 1,773 kinases marked as having active conformations and 1,592 structures in inactive conformations. This dataset is referred to as the KinaseActive dataset in the text. A subset of 514 cyclin-dependent kinases from this set was further analyzed and referred to as the CDKActive dataset.

#### 2.2.2 Shape-mers

Geometricus count vectors were calculated for kinase structures using a *k*-mer size *k* = 20 and a resolution *m* = 2. These were visualized using a t-SNE embedding calculated using the scikit-learn Python library (Pedregosa *et al.*, 2011) with a perplexity of 30 and default parameters.

Shape-mers distinguishing a kinase group *G* were found as those shape-mers which are present in > 95% of the proteins within *G* and whose mean count value within *G* is at least one more than the mean count outside *G*. We visualized distinguishing shape-mers for the STE, AGC, and TK kinase groups using three representative structures, one from each group, with PyMol (DeLano *et al.*, 2002). Each shape-mer can have multiple occurrences across a protein, some of which are shared across groups and do not contribute to the distinguishing nature of the shape-mer. To overcome this, we only visualize occurrences of a group’s shape-mer in the group’s structure which are absent in similar positions across the two structures from the other groups.

Agglomerative clustering was performed on the count vectors, again using the scikit-learn library (Pedregosa *et al.*, 2011), with the Bray-Curtis affinity metric and a distance threshold of 0.63. This threshold was decided using an all vs. all pairwise alignment for 232 kinase structures, up to 40 from each kinase family, from which we took the mean Geometricus similarity score of pairs with an alignment TM-score > 0.95.

#### 2.2.3 Alignment

Subsets of kinase structures were aligned using Caretta-shape with the same parameters as used in benchmarking. For the CK2 alignment, we superposed 292 structures according to the aligning positions and depict each structure as gray lines passing through the *α*-carbon coordinates using Matplotlib (Hunter, 2007). The mean and standard deviation of all coordinates were depicted as a black line and colored circles respectively.

#### 2.2.4 Machine Learning

Gradient Boosting trees were used for machine learning tasks due to their high generalization potential and capability to include missing features as a separate category. These were implemented using the XGBoost Python library (Chen and Guestrin, 2016) with a maximum depth of 5 and remaining default parameters. Kinase active vs. inactive state classification was performed on the KinaseActive and CDKActive datasets with five-fold cross validation. For the KinaseActive dataset consisting of divergent proteins, Geometricus count vectors were used as features. For the CDKActive dataset consisting of the structurally similar cyclin-dependent kinases, aligned moment invariant values were used.

In both cases, feature importance values of each predictor were averaged across cross-validation folds and summed across features. The top 2 predictive shape-mers from the KinaseActive classifier and top 10 predictive residues from the CDKActive classifier are considered in the text. Predictive shape-mer occurrences were mapped back to their corresponding residues. Predictive shape-mer residues and predictive residues from the alignment-based approaches were visualized on representative structures using PyMol (DeLano *et al.*, 2002).

## 3 Results

### 3.1 Fast and accurate multiple structure alignment with rotation-invariant moments

Most machine learning algorithms accept a tabular, fixed dimensional matrix as input, with rows representing individual data points and columns representing features measured across all data points. For proteins sharing high structural similarity this can be accomplished by organizing residue-level features in the order dictated by a multiple structure alignment. Desired properties of this alignment would be high accuracy in terms of structural equivalence of residues, high coverage in order to include as many relevant residue positions as possible instead of just highly conserved positions, and high speed to be able to align and re-align large datasets of proteins in typical parameter selection and validation pipelines. Here we demonstrate that Caretta-shape possesses all three of these properties.

We benchmarked Caretta-shape with the Homstrad (Mizuguchi *et al.*, 1998) and SABMark-superfamily (Van Walle *et al.*, 2004) alignment datasets, and compared against two popular structure aligners, Matt (Menke *et al.*, 2008) and mTM-align (Dong *et al.*, 2017a). Table 2 shows average quality metrics across these datasets and demonstrates that Caretta-shape returns high quality, accurate alignments with high coverage. The pairwise alignment step in Matt and mTM-align is prohibitive, with runtime complexities of *O*(*n*^2^*l*^3^*log*(*l*)) and *O*(*n*^2^*l*^2^) respectively (where *n* is the number of proteins and *l* is the length of the longest protein). mTM-align’s authors mention that 80-90% of their running time is spent in this step (Dong *et al.*, 2017a). Shape-tuple similarity reduces this step to *O*(*n*^2^) in Caretta-shape. The entire Homstrad dataset takes only 4 minutes to align with Caretta-shape, compared to half an hour using the old Caretta algorithm and mTM-align, both of which are 10-15 times faster than Matt (Dong *et al.*, 2017a).

**Table 2:** Average TM-score and percentage of gapless columns across Homstrad and SABmark-superfamily datasets. As the Homstrad dataset also provides reference alignments, “Acc.” shows the number of gapless columns present in the corresponding reference alignment divided by the total number of gapless columns in the reference alignment.

| Aligner | Homstrad |  |  | SABMark-superfamily |  |
| --- | --- | --- | --- | --- | --- |
|  | TM-score | % Gapless | Acc. | TM-score | % Gapless |
| mTM-align | 0.88 | 0.61 | 0.84 | 0.77 | 0.32 |
| Matt | 0.85 | 0.56 | **0.87** | 0.68 | 0.25 |
| Caretta | **0.92** | 0.73 | **0.87** | **0.82** | **0.46** |
| Caretta-shape | **0.92** | **0.74** | **0.87** | 0.81 | 0.45 |

Figure 1 shows the runtime of Caretta-shape on a single thread across synthetic datasets with differing lengths and numbers of proteins. Over a thousand medium-length proteins can be aligned in as little as 20 minutes on a personal computer with a single thread. Further speed improvements such as those employed by multiple sequence alignment algorithms (Sievers *et al.*, 2011) or by the use of graphical processing units (GPUs) could extend Caretta-shape to aligning hundreds of thousands of protein structures in hours; these approaches are left for further exploration.

**Figure 1:**
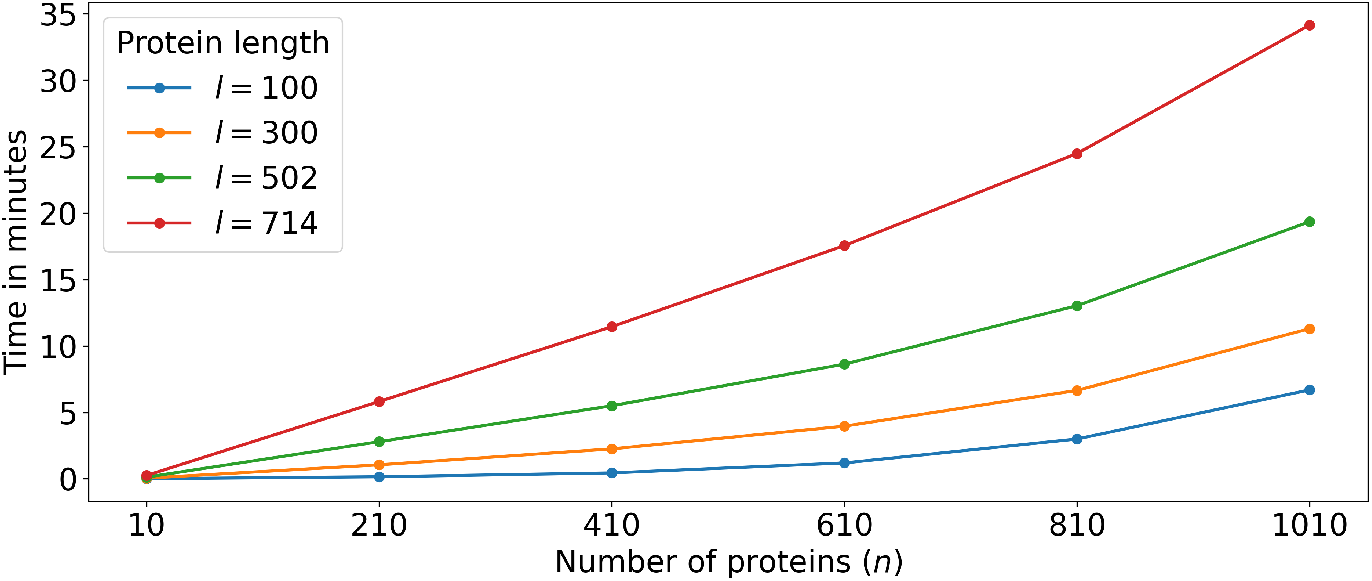
Running time in minutes of Caretta-shape on synthetic datasets with differing number of proteins and proteins with different lengths.

### 3.2 Structure-based exploration of protein kinases

In the past decades, protein kinases have become an alluring target for drug discovery due to the important role they play in key signal transduction pathways. These phosphotransferase enzymes mediate the transfer of the phosphate moeity from high-energy molecules such as ATP to their substrates, and are classified into broad groups based on the substrates they act on. Their popularity has led to a boom in the number of experimentally solved kinase structures with different ligands and inhibitors bound. The kinase–ligand interaction fingerprints and structure database (KLIFS) (Kooistra *et al.*, 2016) now contains 7,746 monomeric structures covering 308 kinases across 8 groups and 3,341 unique ligands.

This superfamily as a whole has divergent protein structures for which only a small 85-residue catalytic segment can successfully be aligned (Van Linden *et al.*, 2014). However, individual kinase families each consisting of up to a thousand structures, share common structural folds that lend well to alignment. With a combination of Caretta-shape and Geometricus we are able to pinpoint differences between kinase groups, align kinase families, and predict conformational change across and within kinase families all in a matter of an hour.

#### 3.2.1 Divergent groups of proteins

Figure 2A shows a t-SNE embedding of Geometricus shape-mer count vectors of all 7,746 kinase monomers in the KinaseAll dataset, colored by the group in which they belong. Clear separation is seen between groups, with smaller clusters visible within each group. These mostly correspond to the kinase families, some of which are labelled in the figure. In Figure 2B, for three kinase groups we look at some shape-mers present in the members of that group and absent in the others. Many of these regions in the structure do not lie in the alignable catalytic stretch and thus would not have been found using alignment-based methods.

**Figure 2:**
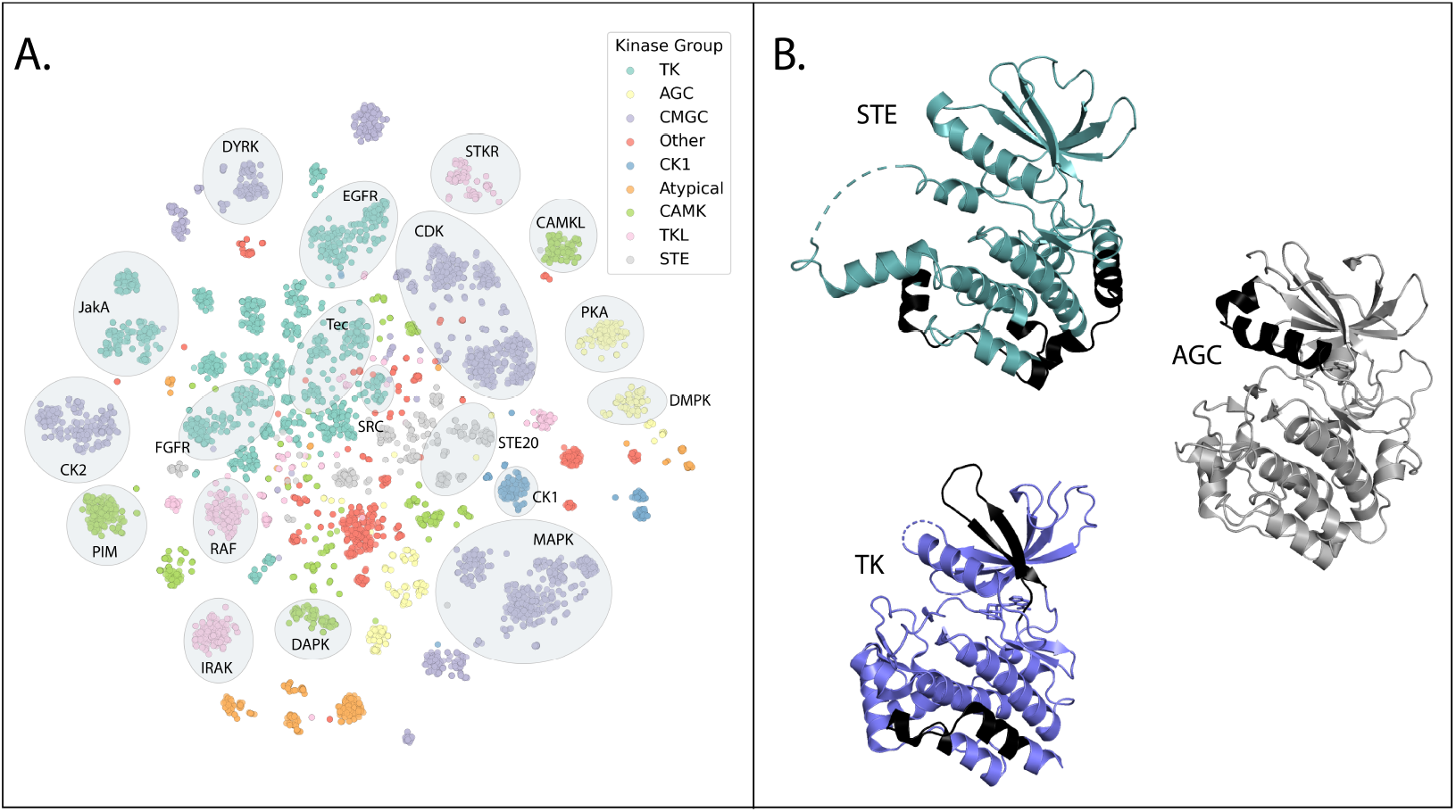
**A**. t-SNE dimensionality reduction of Geometricus shape-mer count vectors for 7,746 kinase monomers, colored by kinase group. 19 kinase families, corresponding to the clusters in Table 3, are labelled. **B**. Shape-mers found in one group and absent in the others, colored black for representative structures of the STE (PDB ID: 4USE), AGC (PDB ID: 3OCB), and TK (PDB ID: 6AAH) kinase groups.

#### 3.2.2 Similar families of proteins

By clustering Geometricus count vectors, we arrive at clusters of proteins displaying high structural similarity which are better suited to alignment. Table 3 shows clusters with > 100 proteins obtained after performing agglomerative clustering with a distance threshold derived from comparison of Geometricus similarity scores with pairwise alignment TM-scores (described in Methods). Each cluster only contains proteins from a single kinase group and many are dominated by a single kinase family (labelled in Figure 2A), demonstrating that Geometricus similarity scores can be used to assign proteins to functional groups when annotations are lacking. We used Caretta-shape to align the proteins within each cluster. The average TM-score of each cluster alignment (reported in Table 3) is very high, confirming their structural similarity. Figure 3 shows the coordinate standard deviations for the CK2 alignment, demonstrating how alignments can be used to assess residue conservation and pinpoint outliers or sub-groups.

**Table 3:** Clusters of kinases obtained from Geometricus count vector clustering, also labelled in Figure 2A. For each cluster we report the average TM-score of its Caretta-shape alignment, the group in which its members belong, and the most frequent family along with the percentage of the family’s occurrence in the cluster (purity).

| No. of proteins | Avg. TM-score | Group | Top family | Purity (%) |
| --- | --- | --- | --- | --- |
| 741 | 0.96 | CMGC | CDK | 82 |
| 661 | 0.96 | CMGC | MAPK | 100 |
| 570 | 0.94 | TK | Tec | 28 |
| 454 | 0.95 | TK | JakA | 59 |
| 346 | 0.95 | TK | FGFR | 61 |
| 292 | 0.99 | CMGC | CK2 | 100 |
| 288 | 0.94 | TK | EGFR | 100 |
| 248 | 0.95 | CMGC | DYRK | 64 |
| 230 | 0.96 | AGC | PKA | 62 |
| 176 | 0.99 | CAMK | PIM | 100 |
| 163 | 0.96 | CAMK | DAPK | 73 |
| 157 | 0.98 | TKL | RAF | 100 |
| 157 | 0.98 | TKL | IRAK | 100 |
| 141 | 0.98 | TKL | STKR | 100 |
| 139 | 0.98 | CAMK | CAMKL | 100 |
| 137 | 0.96 | STE | STE20 | 100 |
| 125 | 0.98 | CK1 | CK1 | 91 |
| 117 | 0.98 | AGC | DMPK | 100 |
| 108 | 0.96 | TK | Src | 66 |

**Figure 3:**
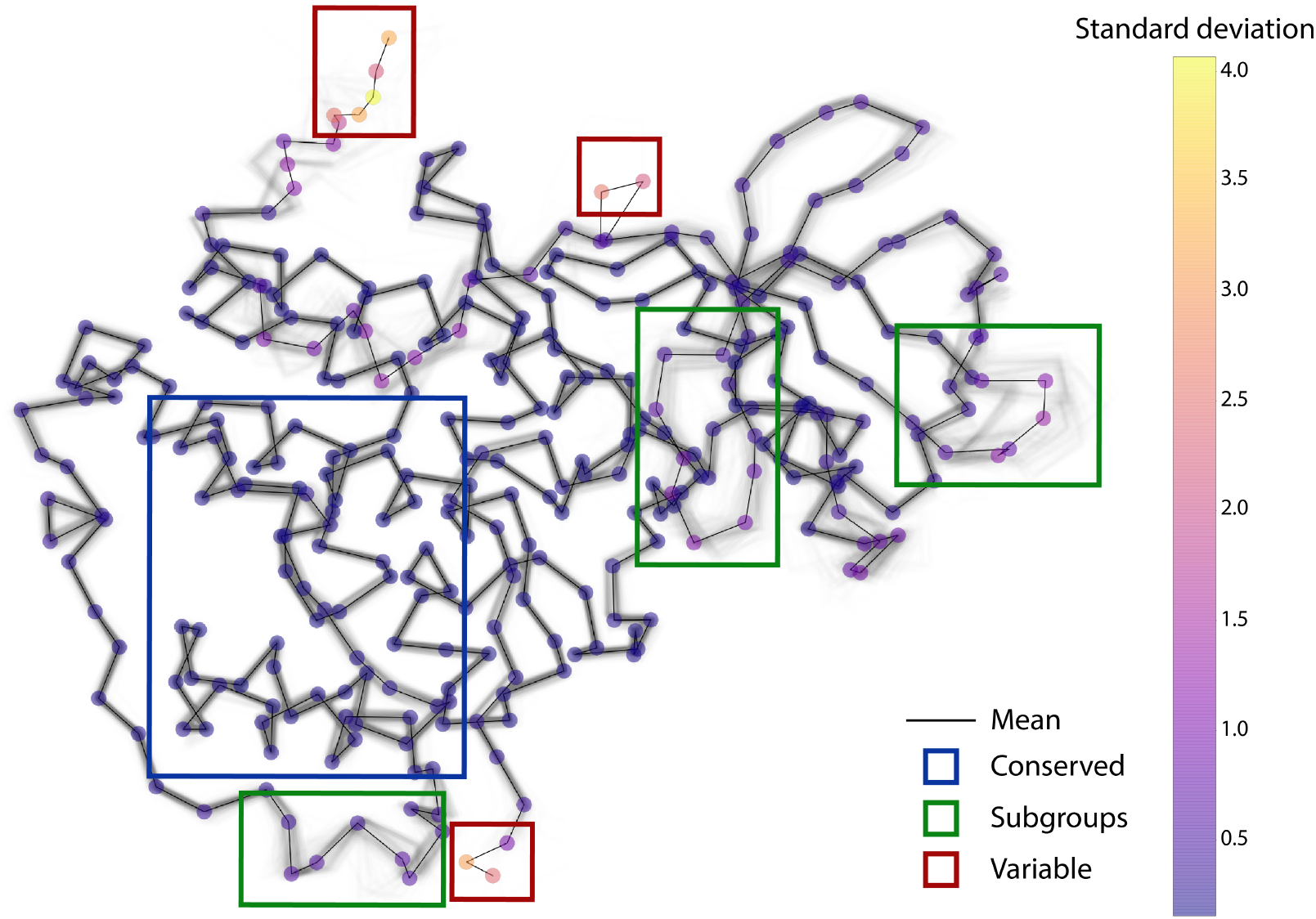
292 CK2 kinase structures superposed according to their Caretta-shape alignment, each depicted as gray lines passing through the *α* carbon coordinates. The mean is depicted as a black line with each residue position colored according to its standard deviation. The blue box marks a well-conserved region, green boxes mark regions showing subgroups where different structures follow different distinct paths (visible in lighter gray), and red boxes mark outlier regions where each protein has highly differing coordinates.

### 3.3 Kinase activity from different perspectives

The protein kinase domain can undergo dramatic conformational changes when reacting to regulatory signals in signaling pathways. These changes are controlled by protein-protein interactions, phosphorylation, and ligand binding (Johnson *et al.*, 1996). Drug discovery efforts often aim to target specific kinase conformations and thus benefit from an understanding of conformational activation across kinases and how this activation differs across the different kinase groups and families.

Using a dataset of 3,365 kinase structures labelled as being in active or inactive conformations (McSkimming *et al.*, 2017), we aim to classify a structure as belonging to one of these two states as well as pinpoint structural elements responsible for the change. We demonstrate how the alignment-free Geometricus is well-suited to tackle such classification problems across diverse proteins, such as those belonging to the different kinase groups, and Caretta-shape alignment allows for zooming into the idiosyncrasies of a single family.

#### 3.3.1 Activation across divergent kinases

To inspect activation across all kinases, we trained a Gradient Boosting classifier on Geometricus embeddings of the KinaseActive dataset. The five-fold cross validation accuracy of this classifier was 96 ± 0.01%. Figure 4A shows the top two predictive shape-mers and their prevalence across active and inactive kinases. These shape-mers are also depicted on an example kinase structure (PDB ID 1E9H). One shape-mer, in dark blue, is localized in the DFG motif which lies in the well-established activation segment (McSkimming *et al.*, 2017). Another, in green, lies in the linker region connecting the activation segment to the *α*F-helix which acts as an organizing hub in the activation process (Kornev *et al.*, 2008). The DFG motif shape-mer is repeated (in light blue) but since Geometricus works with counts we cannot distinguish the true predictive motif using a single structure. More clarity is obtained when looking across multiple structures, as the dark blue occurrence is present in many structures in the active conformational state while the light blue occurrence is not.

**Figure 4:**
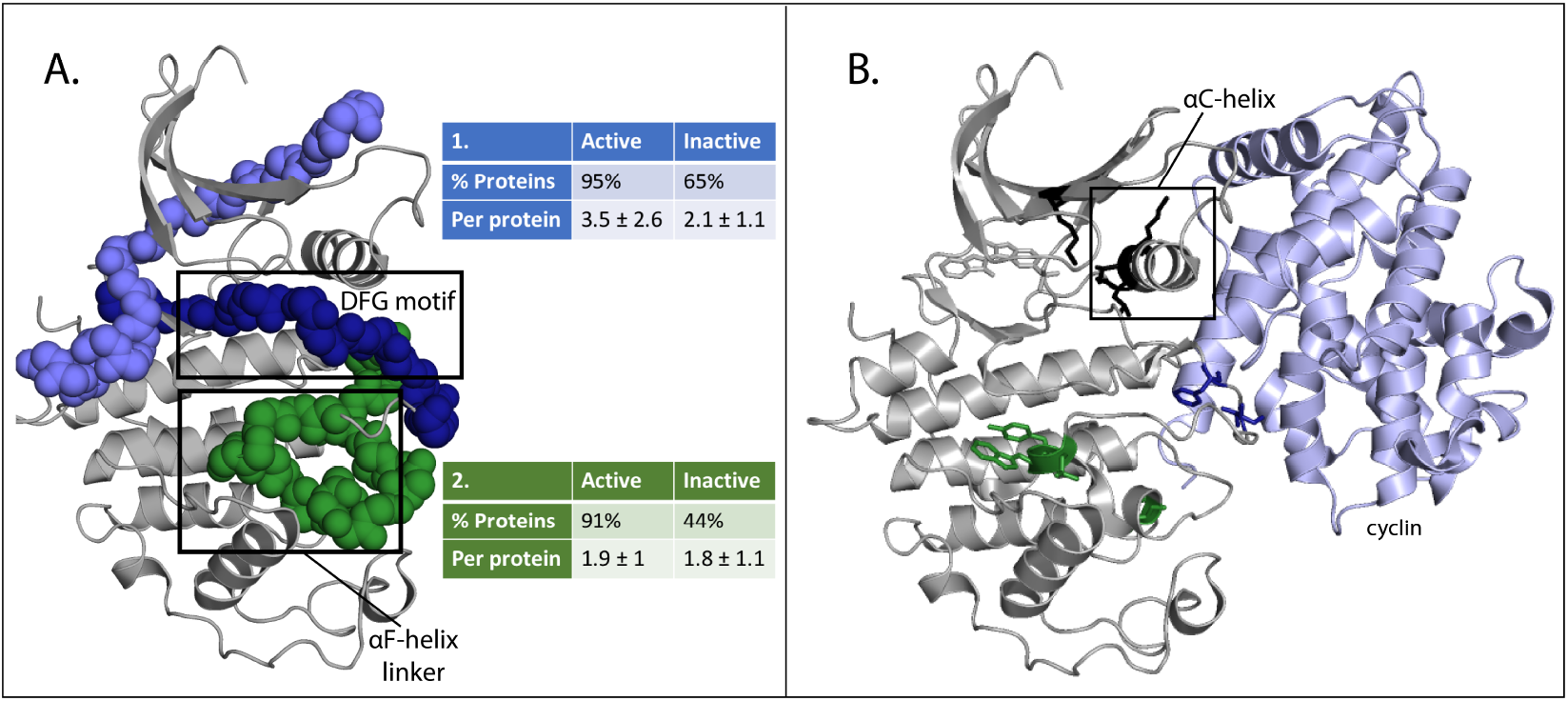
**A**. Occurrences of two shape-mers (shape-mer 1 in dark and light blue, shape-mer 2 in green) found predictive for conformational state classification on a representative kinase structure (PDB ID: 1E9H). The DFG motif and *α*F-helix linker are labeled. The percentage of proteins containing each shape-mer and the average number of times the shape-mer appears per protein across the active and inactive kinases is shown. **B**. Top ten predictive residues for cyclin-dependent kinase conformational state classification shown along with the position of cyclin (light blue). Residues near the DFG motif and *α*F-helix linker are colored blue and green respectively, while the remaining predictive residues are colored black. The *α*C-helix in the CDK-cyclin interface is labelled.

#### 3.3.2 Activation in the cyclin-dependent kinase family

While the Geometricus approach gives us good prediction performance and pinpoints critical structural regions, it misses some structural regions that are specific to certain families. For instance, the cyclin-dependent kinase (CDK) family is dependent on the formation of a CDK-cyclin com-plex. Upon binding, cyclin induces conformational changes in the kinase domain that allow for autophosphorylation of the activation segment to produce a fully active kinase (Russo *et al.*, 1996). Thus, CDKs are further allosterically regulated through cyclin-binding, an aspect not seen in our coarse-grained classifier trained across all kinases. To analyze a specific subfamily such as CDKs, an alignment based approach can be beneficial due to the high structural similarity between proteins and expected residue correspondences. We create a Caretta-shape alignment across 514 CDKs, resulting in an alignment of 399 residues with an average TM-score of 0.96. A Gradient Boosting classifier is trained on the aligned moment invariant values of each CDK, resulting in a very high accuracy of 99%. Figure 4B depicts the top 10 predictive residues. While some residues are again found in the DFG motif and *α*F-helix linker regions, residues in the *α*C-helix which forms part of the cyclin-CDK interface are also found as predictive, indicating that this predictor picks up CDK-specific patterns relevant for kinase conformational change.

## 4 Conclusion

With growing numbers of experimentally solved protein structures and proteins capable of being computationally modeled, structures are seeing increasing use in machine learning applications. To that end we present Caretta-shape, a very fast and accurate multiple structure alignment algorithm based on the concept of rotation-invariant moments, aimed at generating aligned structural features for machine learning.

Depending on the similarity between proteins under study, an alignment-free or alignment-based approach is preferred and each presents its own advantages and insights. We adapt these two approaches to the protein kinase superfamily, which consists of structurally divergent protein groups as well as more similar protein families. We use machine learning to tackle active/inactive conformational state prediction across all kinase families with Geometricus and across the cyclin-dependent kinase family with Caretta-shape alignments. These two approaches lead to the exploration of different aspects of catalytic mechanisms: one aspect explains commonalities within all proteins in this diverse superfamily, and the other zooms in on peculiarities displayed by a single family.

Computational structure modelling is capable of expanding datasets of proteins into the thousands. Once the expensive but automated modelling steps are complete, analyses similar to the ones presented here, both unsupervised and supervised, can be carried out with comparable ease allowing for fast iteration and adaptive exploration of protein biology.

## Broader Impact

The research presented here includes a novel multiple structure alignment algorithm and a demonstration of recently developed algorithms for analysing protein structures with machine learning. Researchers in structural bioinformatics and enzymology may benefit from this work for obtaining structural insights from their data. The ideas discussed also form a fertile basis for more complex algorithms that leverage the increasing amounts of data and recent advances in machine learning and deep learning techniques aimed at such structured, high-dimensional data.

## Notes

https://github.com/TurtleTools/caretta

https://github.com/TurtleTools/geometricus

### Competing Interest Statement

The authors have declared no competing interest.

